# Targeted protein degradation in lysosome utilizing naturally produced bifunctional antibodies with high levels of mannose 6-phosphate glycans

**DOI:** 10.1101/2024.09.03.611037

**Authors:** Andrew C. Hedman, Shou Liu, Jennifer A. Srnak, Riley N. Marcinczyk, Stephanie Do, Linda M. Lyons, Stuart Kornfeld, Hung Do, Lin Liu

**Affiliations:** M6P Therapeutics Inc, St. Louis, MO; Department of Medicine, Washington University School of Medicine, St. Louis, MO

**Author notes:** Correspondence to: Hung Do, PhD, Lin Liu, PhD, Address: 11861 Westline Industrial Dr, Suite 950 St. Louis, MO, 63146. These authors contributed equally to this work (Andrew C. Hedman, Shou Liu).

## Abstract

Novel antibodies have been created for targeted degradation of extracellular and membrane proteins in the lysosome. The mechanism of degradation of target proteins for these antibodies has involved either chemical conjugation of synthetic mannose 6-phosphate (M6P) or engineered bispecific antibodies. Currently, recombinant antibodies cannot be produced with naturally phosphorylated N-glycans. Here, we report the development of a novel platform technology for producing bifunctional therapeutic antibodies with high levels of M6P-bearing glycans directly from producing cells. The antibodies designated as <u>p</u>hosphorylated <u>N</u>-glycosylated peptide <u>c</u>himeric <u>a</u>ntibodies (PNCA) maintain their affinity for antigens with concurrent high affinity binding to cell surface cation-independent mannose-6-phosphate receptors that facilitate internalization and delivery of antibody/antigen complexes to lysosomes for efficient degradation of both target extracellular soluble and membrane proteins. This PNCA approach provides a simple, scalable, and viable approach for producing naturally phosphorylated bifunctional antibodies from production cell lines for targeted protein degradation in lysosomes.

## Introduction

In the past two decades, targeted protein degradation (TPD) has emerged as a novel therapeutic approach for the treatment of many diseases such as cancer, immune diseases, and infections[1–3]. TPD relies on modifications of small molecules or biologics to efficiently remove deleterious proteins through the cellular proteasome or lysosome pathways. The first protein degrader, known as proteolysis-targeting chimera (PROTAC), was reported in 2001[4]. Recently, new technologies such as: LYTAC[5], GalNAc-LYTAC[6], GELYTAC[7], KineTAC[8], Nano-LYTAC[9] and TransTAC[10] have been reported to successfully remove extracellular proteins and/or membrane receptors via lysosome degradation. The mechanism involves modified antibodies which bind to the protein-of-interest (POI) and cell surface internalization receptors. Through efficient receptor mediated endocytosis, the receptor-antibody-POI complex is internalized and ultimately delivered to the lysosome for degradation.

The cation-independent mannose 6-phosphate receptor (CI-MPR) on cell surfaces enables internalization of extracellular lysosomal enzymes and delivery to the lysosome[11]. The CI-MPR binds exogenous lysosomal enzymes bearing the mannose 6-phosphate (M6P) moiety with high binding affinity (nanomolar) at neutral pH. After endocytosis, the CI-MPR/lysosomal enzyme complex is transported to the late endosome/lysosome where the lower pH (pH 4.5 – 5.5) causes a conformation change in the receptor which reduces its binding affinity for M6P and results in the release of lysosomal enzymes. The receptor is then recycled back to the *trans-*Golgi and plasma membrane. The discovery of M6P and the CI-MPR pathway in 1970-80s greatly facilitated the development of recombinant lysosomal enzymes to treat lysosomal storage disorders (LSDs)[12–17]. The formation of the M6P moiety on N-glycans of newly synthesized lysosomal enzymes occurs in *cis*-Golgi via the resident enzyme GlcNAc-1-phosphotransferase (PTase)[18]. The PTase is highly selective for soluble lysosomal enzymes and cannot add M6P to non-lysosomal proteins such as therapeutic antibodies.

In this study, we developed a novel approach which enables natural cellular production of recombinant proteins or antibodies containing high levels of M6P for lysosomal targeting, termed <u>p</u>hosphorylated <u>N</u>-glycosylated peptide <u>c</u>himeric <u>p</u>roteins (PNCPs) or <u>a</u>ntibodies (PNCAs). PNCAs are chimeric recombinant antibodies built on immunoglobulin sequences for antigen recognition and are fused with novel short peptides containing potential N-glycosylation sites at the C-terminus of heavy chain. This novel approach also utilizes a truncated version of PTase, designated as S1S3 PTase, that was strategically engineered to produce a pre-activated holoenzyme that has approximately 20-fold higher phosphate transferring activity than the wild-type PTase in cells[19, 20]. The data in this study show that four PNCAs produced from suspension ExpiCHO cells are highly effective for binding cell surface CI-MPRs for internalization and delivering extracellular soluble target proteins and membrane proteins to lysosomes for subsequent degradation.

## Results

### Generation of new glycoproteins with high levels of mannose 6-phosphate (M6P)

To generate high M6P levels on non-lysosomal proteins to enable lysosomal targeting, we created short novel peptide sequences containing multiple potential N-glycosylation sites (N-X-T/S) to fuse with non-lysosomal proteins. The peptide containing 6 potential N-glycan sites was fused to the C-terminus of enhanced green fluorescent protein (GFP) and the Ig Kappa signal sequence was utilized to express a secreted, glycosylated GFP chimeric protein, designated as an <u>N</u>-glycosylated peptide <u>c</u>himeric <u>p</u>rotein-GFP (NCP-GFP). Figure 1A shows a schematic depiction for producing NCP-GFP or <u>p</u>hosphorylated <u>N</u>-glycosylated <u>c</u>himeric GFP <u>p</u>rotein (PNCP-GFP) with S1S3 PTase co-expression in ExpiCHO cells. To determine the types of N-glycans present on the NCP-GFP and PNCP-GFP proteins, conditioned media were treated with Peptide-N-Glycosidase F (PNGase-F) which cleaves off all N-glycans or Endoglycosidase H (Endo-H) which only cleaves high mannose type glycans and analyzed by western blotting using anti-GFP antibody. Untreated NCP-GFP and PNCP-GFP proteins run as larger protein species (with apparent major molecular weights above 43 kDa) while PNGase-F treatment removes all N-glycans resulting in a significant reduction in the apparent molecular weight that is observed slightly above 34 kDa (Fig. 1B). Only PNCP-GFP was sensitive to Endo-H treatment, indicating that only PNCP-GFP contains high mannose type N-glycans which are the requisite glycan structures for potential phosphorylated N-glycans. To confirm that PNCP-GFP contains M6P, the NCP-GFP or PNCP-GFP samples were added to a CI-MPR coated plate to detect bound GFP by green fluorescence. Fig. 1C panel shows that only PNCP-GFP binds to CI-MPR as evidenced by increased GFP fluorescence. Both NCP-GFP and PNCP-GFP samples were shown to have similar total GFP fluorescence signals in conditioned medium that were applied to the CI-MPR binding assay (Fig. 1D). These results demonstrate that the N-glycosylated peptide enabled production of GFP with phosphorylated N-glycans when co-expressed with the S1S3 PTase.

**Figure 1.**
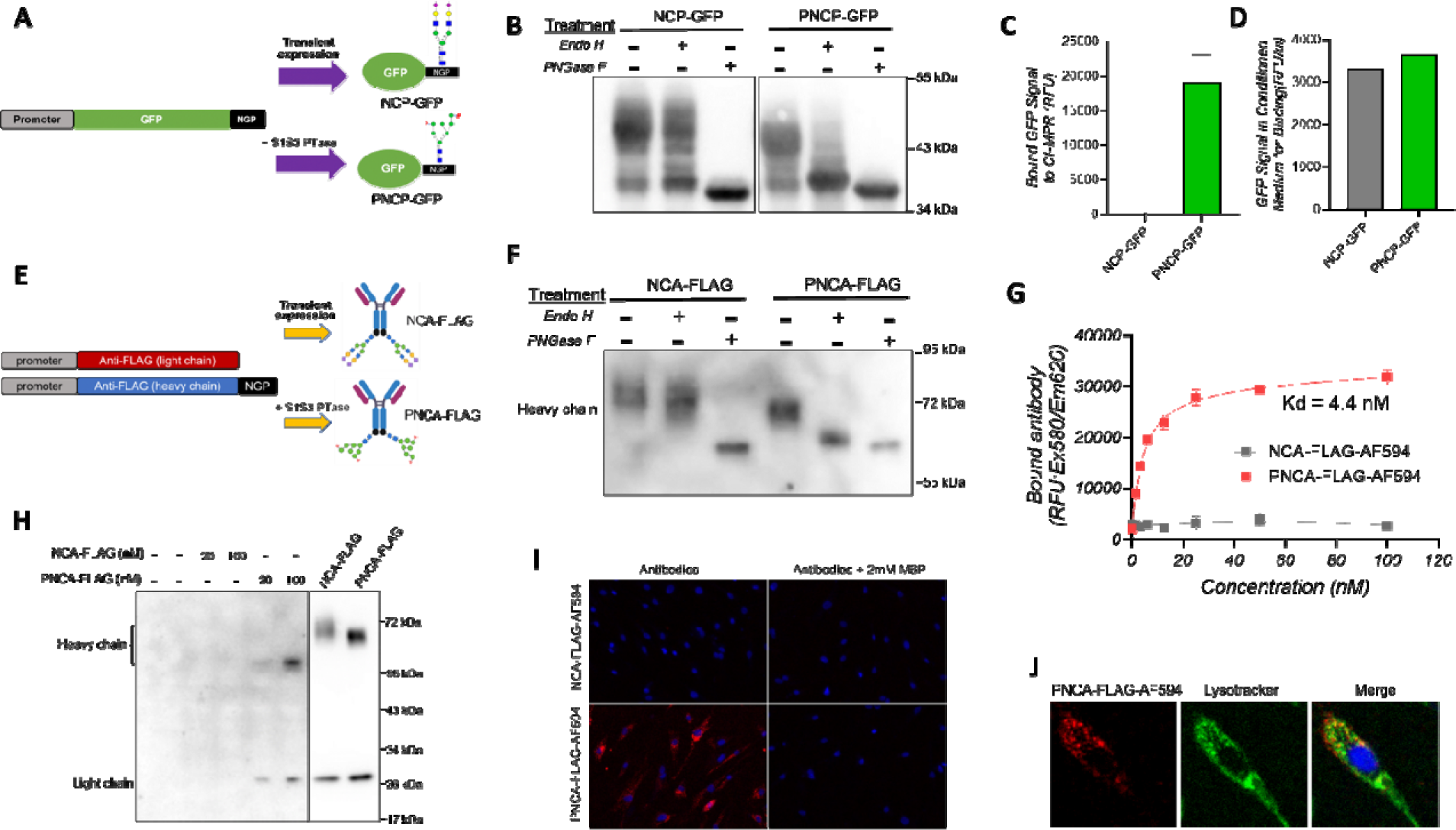
Generation of new N-glycosylated peptide fusion proteins and antibodies with high levels of mannose 6-phosphate (M6P). **A**. Schematic showing production of NCP-GFP and PNCP-GFP in CHO Cells. **B**. Conditioned medium containing NCP-GFP or PNCP-GFP was treated with Endo-H or PNGase-F and samples were examined by western blot to detect the molecular weight shift. **C**. NCP-GFP and PNCP-GFP protein from (B) wa examined for CI-MPR binding in the 96-well plate-based assay. Binding was measured by GFP fluorescence signal. N=3. **D**. GFP fluorescence in the produced conditioned medium used for CI-MPR binding assay. **E**. Schematic grap shows the production of NCA-FLAG and PNCA-FLAG in CHO cells. **F**. Western blotting to examine the NCA-FLAG and PNCA-FLAG heavy chain N-glycosylation after Endo-H and PNGase-F treatment. **G**. CI-MPR plate binding assay to determine the binding of NCA-FLAG and PNCA-FLAG toward CI-MPR. To visualize the binding signal, Alexa Fluor 594 was conjugated to NCA-Flag (gray) or PNCA-Flag (red) for the assay. N=3. **H**. Internalization of NCA-FLAG or PNCA-FLAG in human fibroblast cells treated with at 20 or 100 nM each antibody for 24 hours. Purified antibodies were loaded as loading control. **I**. Alexa Fluor 594 conjugated NCA-GFP or PNCA-GFP were incubated with human fibroblast cells with and without 2 mM free Mannose 6-phosphate in the media for 6 hours. The antibody uptake was examined by Alexa Fluor 594 fluorescence under fluorescent microscopy. Nuclei were stained with Hoechst (blue). **J**. Internalized PNCA-FLAG (red) co-localized with lysotracker (green) in fibroblast cells. Data are represented as mea values; error bars represent the standard deviation of biological replicates.

Next, we tested whether a chimeric antibody containing a C-terminal fusion with the N-glycosylated peptide could be produced with naturally phosphorylated N-glycans from CHO cells. A schematic (Fig.1E) details the construct design for expression of mouse anti-FLAG M2 light chain under CMV promoter and the heavy chain fused with the N-glycosylated peptide under the EF-1α promoter. <u>N</u>-glycosylated peptide <u>c</u>himeric <u>a</u>ntibodies (NCAs) were produced from transiently transfected ExpiCHO cells and purified by protein G. NCA-FLAG (produced without the S1S3 PTase) or PNCA-FLAG (produced with the S1S3 PTase) samples were treated with Endo-H or PNGase-F and the antibody heavy chain molecular weight was examined by SDS-PAGE/western blotting (Fig.1F). NCA-FLAG was sensitive to PNGase-F treatment, but not Endo-H treatment, indicating the presence of only complex type N-glycans. By contrast, the apparent molecular weight of PNCA-FLAG was significantly decreased by both Endo-H and PNGase-F treatment confirming the presence of high mannose type glycans. Next, purified NCA-FLAG and PNCA-FLAG samples were conjugated with Alexa-Fluor 594 (AF594). A concentration dependent increase in AF594 fluorescence signal that was saturated at the higher protein concentrations was observed in AF594 labeled PNCA- FLAG (PNCA-FLAG-AF594) with an estimated binding affinity (Kd) of approximately 4 nM while NCA-FLAG did not bind to CI-MPR at all (Fig.1G & Extended Data Fig.1A). CI-MPR affinity chromatography revealed that more than 90% of PNCA-FLAG produced binds to CI-MPR (Extended Data Fig.1 B&C) as compared to no binding for NCA-FLAG.

To determine if the phosphorylated N-Glycans on PNCA-FLAG can enhance its internalization into cells and delivery to lysosomes, wild-type human fibroblasts were incubated with 20 and 100 nM NCA-FLAG or PNCA-FLAG for 6 hours. Only the phosphorylated PNCA-FLAG was detected in cells by SDS-PAGE/western blotting (Fig.1H). Cellular uptake was also examined by incubating AF594 conjugated NCA- FLAG or PNCA-FLAG in human fibroblasts and analyzed by fluorescence microscopy. Images shown in Fig.1I demonstrate strong intracellular AF594 signal with PNCA- FLAG-AF594, but not with NCA-FLAG-AF594. The uptake was blocked by co-incubation with 2 mM free M6P, indicating that the uptake of PNCA-FLAG-AF594 is mediated by M6P through the CI-MPR pathway. In addition, lysosomal localization of PNCA-FLAG-AF594 (red) was evaluated by co-staining with lysotracker (green), an established marker of the lysosome. The yellow signal (Fig. 1J) in the merged panel represents overlapping red and green fluorescence signal that indicates colocalization of PNCA-FLAG and lysosomes, confirming the delivery of the PNCA antibody to the lysosome. These results indicate that the PNCA produced with S1S3 PTase co-expression contains M6P-bearing N-glycans, has high binding affinity to CI-MPR and can be efficiently internalized and delivered to the lysosome.

### PNCA-FLAG binds to antigen and antibody/antigen complex are internalized and delivered to lysosome through CI-MPR pathway

While PNCA-FLAG was shown to contain M6P-containing N-glycans, these carbohydrate structures do not alter its antigen recognition function as tested by western blot (Extended Data Fig.2). To assess whether PNCA-FLAG can capture antigen and concurrently bind to cell surface CI-MPR, a soluble FLAG-tagged mCherry (FLAG-mCherry) was generated from ExpiCHO cells and CI-MPR plate binding assay was performed by incubating the FLAG-mCherry with either NCA-FLAG or PNCA-FLAG in CI-MPR coated plates (Fig.2A). Fig.2B shows the mCherry fluorescence signal is concentration-dependent and saturable with PNCA-FLAG only, indicating that the PNCA-FLAG can simultaneously bind to both its antigen (FLAG-mCherry) and CI-MPR. Next, we evaluated the antibodies for facilitating cellular uptake of extracellular FLAG- mCherry using HepG2 cells. Cells were incubated overnight with FLAG-mCherry in the presence of 20 nM NCA-FLAG or PNCA-FLAG antibodies and mCherry fluorescence was examined in cells. mCherry signal was detected only in cells treated with PNCA- FLAG but not with NCA-FLAG (Fig.2C). The uptake of FLAG-mCherry into cells via PNCA-FLAG was inhibited by the addition of 2 mM M6P. To better quantify the relative amounts of extracellular antigen that was internalized, we created a construct to express FLAG tagged HexM enzyme, a modified version of the lysosomal enzyme β- hexosaminidase with enhanced stability and ability to homodimerize[21]. As has been shown, overexpression of HexM in cells produces the enzyme with a minimal amount of M6P[22]. FLAG-HexM was produced in ExpiCHO cells, and the conditioned cell culture medium was then co-incubated with 20 nM NCA-FLAG or PNCA-FLAG for 4 hours to assess the amount of internalized antigen. The relative amounts of internalized FLAG- HexM in five different cell models (HEK293T, Tay-Sachs patient fibroblast, Daoy, HepG2, and MDA-MB-231) was determined by HexM enzyme activity in cell lysates (Fig. 2D-H). ′FLAG-HexM alone is internalized at low background levels. Co-incubation with NCA-FLAG did not further increase the HexM activity in cells. In contrast, PNCA- FLAG significantly increased FLAG-HexM internalization by 2.5- to 9-fold higher than NCA-FLAG or FLAG-HexM alone groups as measured by HexM activity in cell lysates. The internalization of FLAG-HexM was inhibited by co-incubation with 2 mM free M6P in the culture medium. To further demonstrate lysosomal targeting was mediated by M6P via the CI-MPR pathway, CI-MPR knockout cells (CI-MPR^-/-^) cells were generated by CRISPR-Cas9 in HEK293T cells (Fig.2I). Only background HexM enzyme activity was detected in each group (Fig.2J), demonstrating that the removal of CI-MPR completely abolished the uptake of FLAG-HexM mediated by PNCA-FLAG. This data indicates that only the antibodies with phosphorylated N-glycans (PNCAs) can internalize extracellular antigens into cells through the CI-MPR pathway.

**Figure 2.**
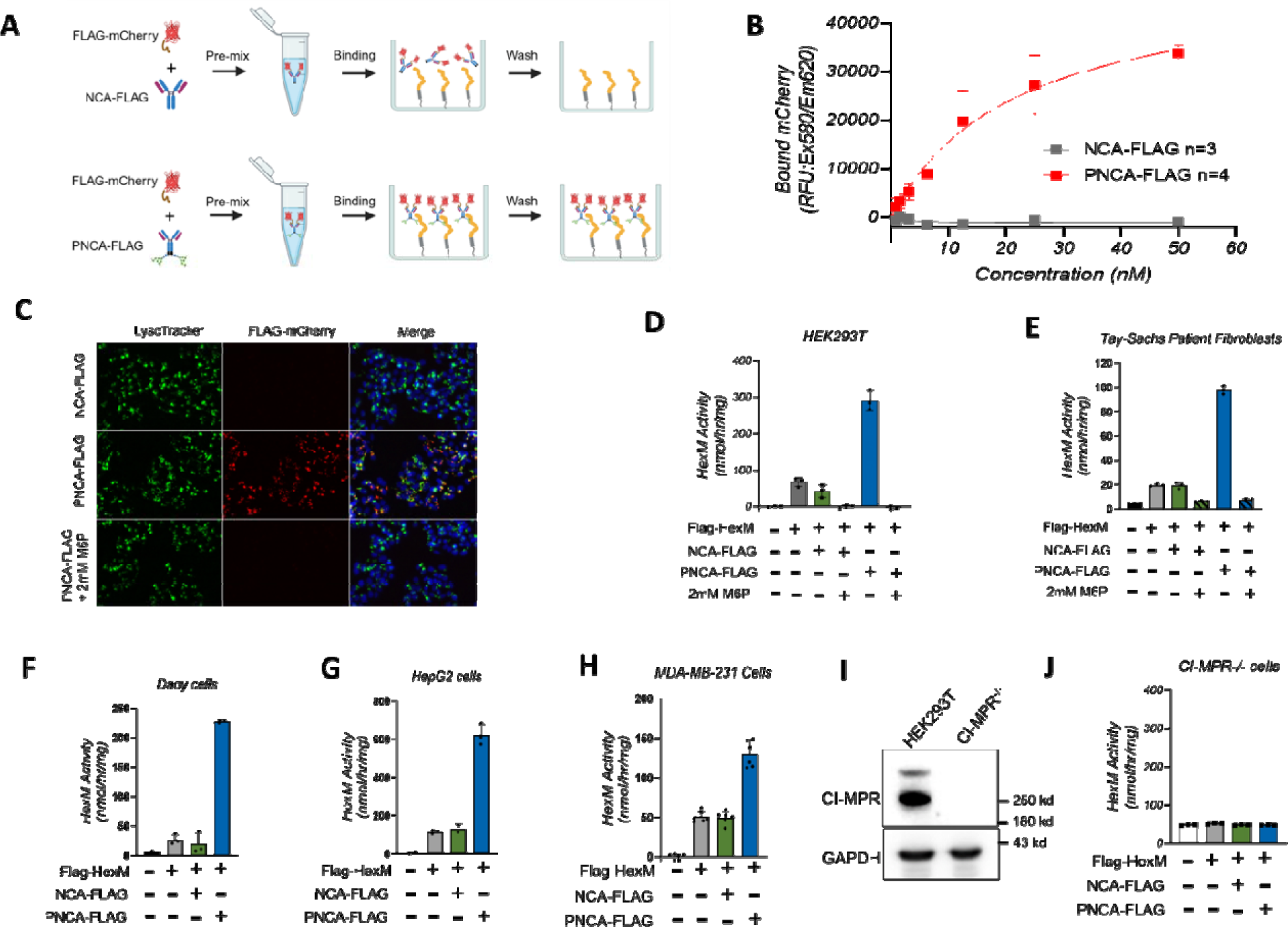
PNCA-FLAG binds to antigen and is internalized and delivered to lysosome with antigen through CI-MPR pathway. **A**. Schematic showing the procedure for detection of FLAG-mCherry and NCA-FLAG or PNCA-FLAG complexes binding toward CI-MPR. **B**. Bound mCherry fluorescence from (A) for NCA-FLAG (gray) or PNCA-FLAG (red). N=3 or 4. **C**. HepG2 cells were incubated with FLAG-mCherry and NCA-FLAG or PNCA-FLAG overnight with or without 2 mM M6P and cells were examined by fluorescent microscopy to detect mCherry signal (red). Lysosomes were stained with LysoTracker (green) and Hoechst (blue) for nuclei. **D-H**. Cell lines (D. HEK293T, E. Tay-Sachs Patient Fibroblasts, F. Daoy, G. HepG2, or H. MDA-MB-231) were incubated with FLAG-HexM enzyme with NCA-FLAG or PNCA-FLAG or with addition of 2 mM M6P. HexM internalization was determined by performing a HexM activity assay and activity graphed – HexM alone (gray), NCA-FLAG with HexM (Green), PNCA-FLAG with HexM (Blue), M6P added (black striped bars). Background HexM activity in untreated group wa subtracted. Dot represents each independent experiment. N=3-6. **I**. HEK293T CI-MPR^-/-^ cells were generated by CRISPR-Cas9. Western blot shows CI-MPR deficiency in the cells. **J**. Graphs depict internalized HexM activity in CI-MPR^-/-^ cells. Dot represents each independent experiment. N=3. Data are represented as mean values; error bars represent the standard deviation of biological replicates.

### PNCA-TNF**α** enables internalization and degradation of extracellular TNF**α** in the lysosome

Neutralizing soluble proinflammatory cytokines using monoclonal antibodies as inhibitors has been utilized therapeutically for decades[23]. For example, Adalimumab works by binding and blocking the tumor necrosis factor-alpha (TNFα) interaction with its receptors[24]. To test if the PNCA technology can be applied to a therapeutic humanized antibody for lysosomal degradation of soluble cytokines, we constructed a <u>N</u>-glycosylated <u>c</u>himeric anti-TNFα <u>a</u>ntibody (NCA-TNFα) comprised of the human IgG1 Fc region fused with the N-glycosylated peptide at the C-terminus of the heavy chain of adalimumab sequence. PNCA-TNFα was generated by co-expression of the chimeric antibody with S1S3 PTase in ExpiCHO cells. In addition, the regular human adalimumab antibody (Ab-TNFα) was generated as a control (Schematic in Fig.3A & Extended Data Fig.3A). Endo-H and PNGase-F analysis were performed in the antibodies. Normal human IgG1 contains a single N-glycan in the heavy chain (N297)[25], such that a small shift was observed with PNGase-F treatment in Ab-TNFα. However, a large molecular weight shift following Endo-H and PNGase-F treatment was observed for PNCA-TNFα confirming the presence of newly introduced high mannose type N-glycans (Fig.3B). Around 85% of the total PNCA-TNFα antibody bound to the CI-MPR receptor by CI- MPR affinity chromatography (Fig.3C&D) with an estimated Kd ∼7 nM on the plate binding assay (Extended Data Fig.3B) using AF594 conjugated PNCA-TNFα. No binding was detected in Ab-TNFα. PNCA-TNFα was shown to facilitate efficient cellular uptake in human fibroblasts as evidenced by the strong intracellular fluorescence signal in cells (Extended Data Fig.3C), while no binding or internalization were observed for Ab-TNFα which lacks M6P.

**Figure 3.**
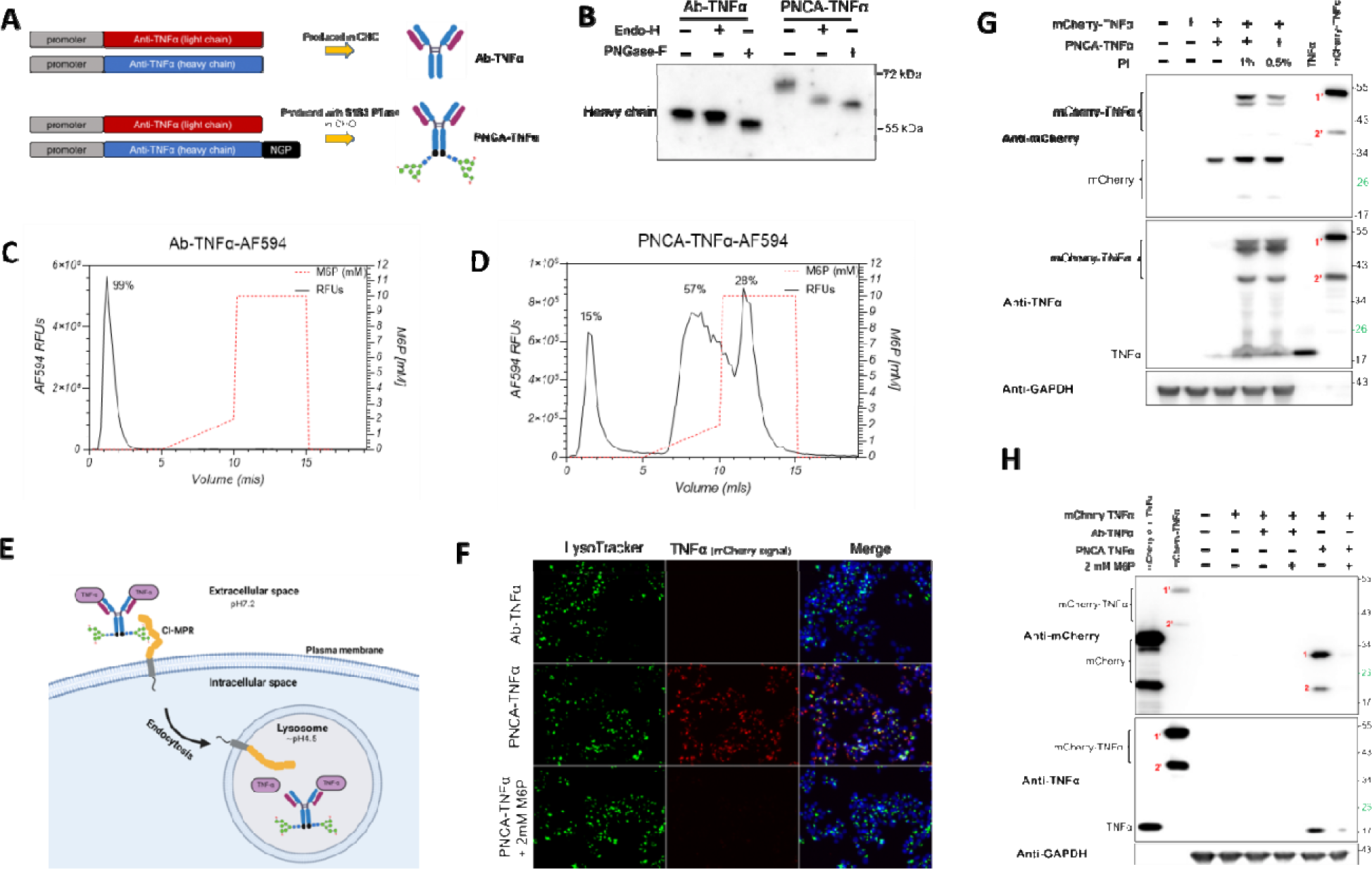
PNCA-TNFα enables internalization and degradation of extracellular TNFα in the lysosome. **A**. Schematic demonstrating production of Ab-TNFα and PNCA-TNFα in CHO cells. **B**. Western blotting to examine the Ab-TNFα and PNCA-TNFα heavy chain N-glycosylation after Endo-H and PNGase-F treatment. **C-D**. CI-MPR affinity chromatograph to determine the binding of Ab-TNFα and PNCA-TNFα. To visualize the binding signal, Alexa Fluor 594 conjugated to Ab-TNFα (C) or PNCA-TNFα (D) was used for the assay. Red dash line indicates the concentration of free M6P for antibody elution from the CI-MPR column. **E**. Schematic depicting internalization and degradation of extracellular TNFα b PNCA-TNFα through CI-MPR to the lysosome. **F**. HepG2 cells were incubated with mCherry-TNFα and Ab-TNFα or PNCA-TNFα overnight with or without 2 mM M6P and cells were examined by fluorescent microscopy to detect mCherry-TNFα signal (red). Lysosomes were stained with LysoTracker (green) and Hoechst for Nuclei (blue). **G**. Internalized mCherry-TNFα protein was analyzed by western blot for cell lysates using antibodies to detect mCherry, TNFα, and GAPDH as a loading control. **H**. Western blot analysis for internalized mCherry-TNFα with the addition of 1% or 0.5% of lysosomal protease inhibitors (PI). Antibodies are used to detect mCherry, TNFα and GAPDH, as a loading control.

Next, antibody mediated TNFα internalization and degradation were examined in cellular models and is schematically depicted in Fig.3E. To monitor TNFα internalization, a fusion protein containing mCherry at the N-terminus of soluble human TNFα (mCherry-TNFα) was expressed, purified and utilized in cellular uptake studies (Extended Data Fig.4). HepG2 cells were incubated with mCherry-TNFα with either Ab- TNFα or PNCA-TNFα in the presence or absence of 2 mM M6P. Internalization of TNFα (mCherry-TNFα fluorescence signal) was only observed with PNCA-TNFα but not with the non-phosphorylated Ab-TNFα (Fig.3F). This uptake was inhibited by the addition of 2 mM M6P. Cellular uptake of mCherry-TNFα was also assessed by SDS- PAGE/western blot analysis using anti-mCherry (top) or anti-TNFα (middle) antibodies (Fig.3G). mCherry-TNFα uptake was observed only with PNCA-TNFα as detected by both anti-mCherry antibody and anti-TNFα antibody. Protease inhibitors are useful tools to confirm the presence of internalized target antigens in cells since they have been shown to inhibit protein degradation in the endosome/lysosome when applied to cells during cell culture[26]. To determine whether the internalized mCherry-TNFα can be degraded in lysosomes, HepG2 cells were treated with mCherry-TNFα alone or with PNCA-TNFα in the presence or absence of protease inhibitors and analyzed by SDS-PAGE/western blotting as shown in Fig.3H. In the absence of protease inhibitors, only low amounts of mCherry and TNFα protein were detected while the full-length mCherry- TNFα and higher amounts of mCherry and TNFα were detected with the inclusion of protease inhibitors. Similar results were observed with recombinant human TNFα as shown in Extended Data Fig.5. Our data indicates that extracellular TNFα can be efficiently internalized into cells via PNCA-TNFα and subsequently degraded in the lysosome. These data show PNCA human antibodies retain their high affinity interactions with their soluble antigens (shown here for TNFα) and the phosphorylated antibody-antigen complex can be internalized by the CI-MPR pathway and delivered to the lysosome for subsequent degradation.

### Native N-glycans on antibodies are poorly phosphorylated

We sought to understand whether the S1S3 PTase can increase M6P levels on the native N-glycans of antibodies without fusion to the N-glycosylated peptide. It should be noted that the native N-glycans on antibodies cannot be phosphorylated when produced in cells with endogenous wild-type PTase as observed by empirical testing of multiple antibodies. Ab-TNFα antibody sequences were expressed with the S1S3 PTase co-expression in ExpiCHO cells, designated as Ab-TNFα-S (Extended Data Fig.6A). Endo-H treatment of Ab-TNFα-S was shown to shift the heavy chain of antibody produced with the S1S3 PTase to a lower apparent molecular weight (Extended Data Fig.6B). The antibody was further conjugated with AF594 and characterized on CI-MPR affinity chromatography. Only a small proportion (19%) of the Ab-TNFα-S antibody binds to the CI-MPR (Extended Data Fig.6C) which is much lower than that observed for PNCA- TNFα (∼85%, Fig. 3D). These data suggest the native N-glycans in the human IgG1 are not sufficient for the generation of M6P even with S1S3 PTase co-expression. Thus, both the N-glycosylated peptide and S1S3 PTase co-expression are required for efficient phosphorylation of N-glycans on recombinant antibodies.

### PNCA antibodies enable degradation of target cell surface transmembrane receptors

Programmed death ligand 1 (PD-L1) is an established biomarker that is over-expressed on the cell surface of tumor cells[27]. Therapeutic antibodies toward PD-L1 have been applied to block the interaction to its receptor (PD1) for cancer treatment. To determine whether PNCA can mediate targeted lysosomal degradation of PD-L1 in cells, anti-PD- L1 antibody – Atezolizumab variable sequence was incorporated into the human IgG1 fusion construct containing our N-glycosylated peptide design. The <u>p</u>hosphorylated <u>N</u>- glycosylated peptide <u>c</u>himeric PD-L1 <u>a</u>ntibody (PNCA-PD-L1) was produced with co-expression of S1S3 PTase and purified. A human IgG1 with Atezolizumab variable domain sequences served as a non-phosphorylated control antibody (Ab-PD-L1) (Extended Data Fig.7A). Western blotting analysis after Endo-H treatment indicates that PNCA-PD-L1 contains high mannose-type N-glycans as evidenced by the significant reduction in the apparent molecular weight of the heavy chain post-treatment (Extended Data Fig.7B). CI-MPR affinity chromatography using the AF594-conjugated antibody shows ∼80% of the PNCA-PD-L1 binds to CI-MPR whereas the non-phosphorylated Ab- PD-L1 antibody does not bind at all (Extended Data Fig.7C&D). A proposed mechanism for PNCA-PD-L1 mediated lysosomal targeting and degradation of PD-L1 in cells is depicted in Fig.4A. PNCA-PD-L1 was shown to be efficiently internalized into wild-type human fibroblasts as detected by the intracellular AF954 fluorescence signal and delivered to lysosomes as evidenced by co-localization with lysotracker (green) in the cells (Fig.4B). To evaluate the impact of PNCA-PD-L1 on PD-L1 levels, Daoy (medulloblastoma tumor) or A431 (epidermoid carcinoma) cells were incubated with 1 to 20 nM Ab-PD-L1 or PNCA-PD-L1 for 24 hours followed by analysis of residual PD-L1 protein levels by SDS-PAGE/western blotting. The PNCA-PD-L1 antibody was shown to facilitate near total elimination of PD-L1 in Daoy (Fig.4C) and A431 (Fig.4D) cells whereas the non-phosphorylated Ab-PD-L1 antibody had a modest effect on PD-L1 levels under identical experimental conditions.

**Figure 4.**
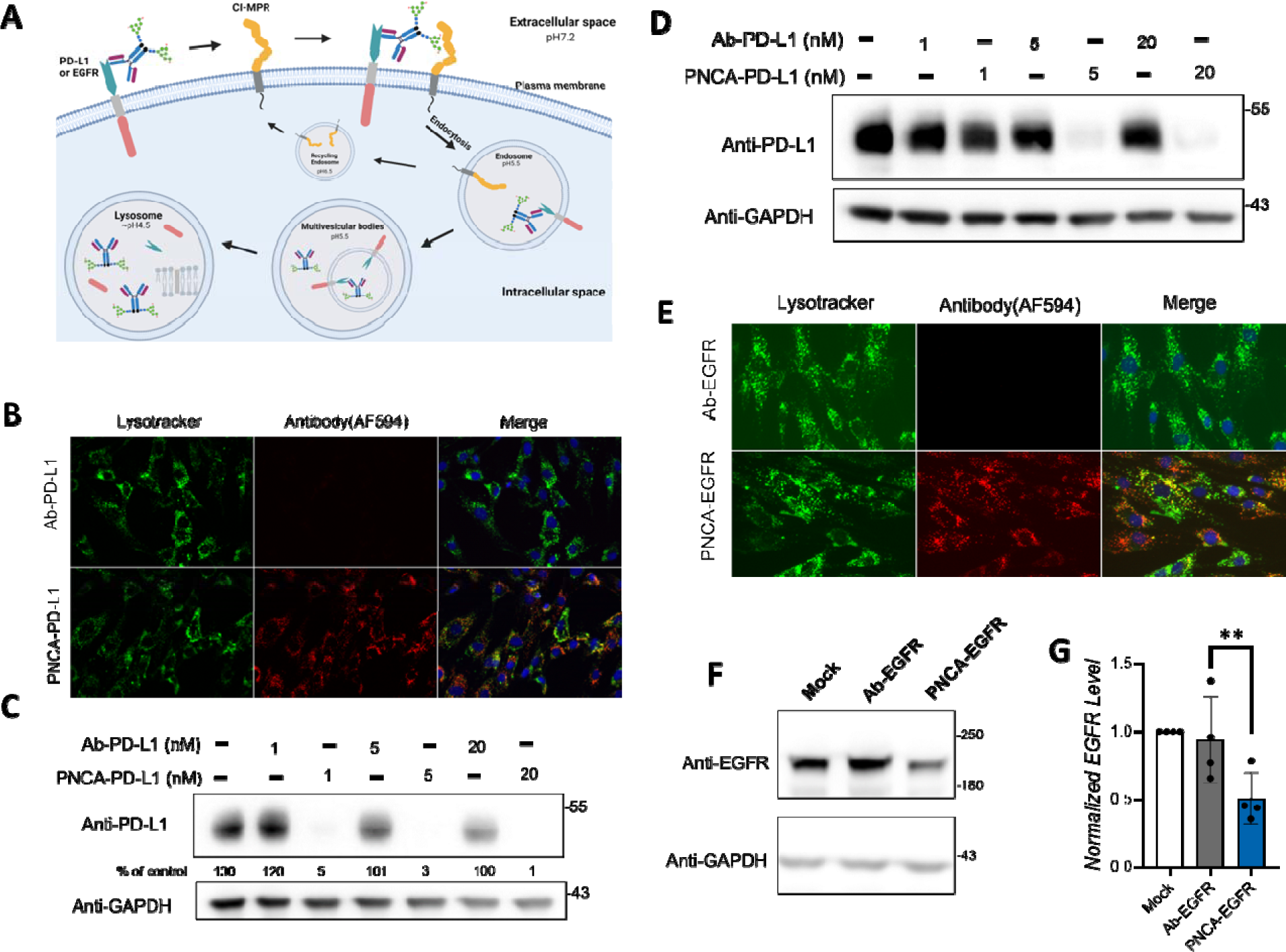
PNCA antibodies enable degradation of target cell surface transmembrane receptors. **A**. Schematic depicting internalization of cell surface PD-L1 or EGFR by PNCA antibodies and CI-MPR to mediate receptor internalization and lysosomal degradation. **B**. Internalization of Ab-PD-L1 and PNCA-PD-L1 in fibroblast cells. AF594 labeled antibodies were incubated with the cells and co-stained with LysoTracker (green) and Hoechst (blue). **C**. Daoy cells were incubated alone or with 1, 5 or 20 nM Ab-PD-L1 or PNCA-PD-L1 for 24 h. Western blot was performed on cell lysates to detect total PD-L1 or GAPDH, loading control. The relative amounts of PD-L1 were normalized to GAPDH in each cell lysate sample (expressed as % of control) to estimate the residual amount of PD-L1 after treatment. **D**. A431 cells were treated similarly as (C). Westen blot to check the total PD-L1 level. **E**. Human fibroblast cells were incubated with AF594 conjugated purified Ab-EGFR or PNCA-EGFR. Antibody uptake is indicated by red signal. Lysosomes were stained by incubation with lysotracker (green), and nuclei were stained using Hoechst (blue). **F**. Daoy were treated using 20 nM of Ab-EGFR antibody or PNCA-EGFR for 24 h. Cell lysates were examined by Western blot for total EGFR protein or GAPDH. **G**. Quantification of (F). Dot represents each independent experiment. N=4. Data are represented as mean values; error bars represent the standard deviation of biological replicates. Significance testing was performed by paired two-tailed t-tests.

The epidermal growth factor receptor (EGFR) regulates epithelial tissue development and homeostasis and is expressed in a variety of human tumors[28]. The commercial anti-EGFR (Cetuximab) antibody sequence was utilized to create phosphorylated anti- EGFR antibodies. Regular non-phosphorylated anti-EGFR antibody (Ab-EGFR) and phosphorylated anti-EGFR antibody (PNCA-EGFR) were transiently expressed from ExpiCHO cells and purified (Extended Data Fig.8A&B). PNCA-EGFR was shown to contain Endo-H sensitive N-glycans (indicative of high mannose-type N-glycans), binds to CI-MPR affinity chromatography at similarly high levels (82%) as other PNCAs, has high binding affinity (Kd∼10 nM) in CI-MPR plate binding assays, and was efficiently internalized into human fibroblast and HepG2 cells (Extended Data Fig.8C-H &Fig.4E). The PNCA-EGFR antibody significantly reduced the total EGFR level in Daoy cells, whereas the non-phosphorylated Ab-EGFR antibody had no effect (Fig. 4F&G). These results indicate that the PNCA antibodies with M6P-containing N-glycans can bind to both the target transmembrane antigen and CI-MPR on the cell surface to facilitate endocytosis and delivery of target cell surface receptors to lysosomes for degradation.

## Discussion

Our data show that the <u>p</u>hosphorylated <u>N</u>-glycosylated peptide <u>c</u>himeric <u>p</u>rotein/<u>a</u>ntibody (PNCP/PNCA) produced with S1S3 PTase co-expression in CHO cells have high binding affinity toward CI-MPR and enable robust lysosome targeting and degradation of both extracellular soluble and cell surface membrane proteins. Unlike typical therapeutic antibodies that bind to target antigens to sterically block their interactions with other proteins, our data indicate that PNCA antibodies not only bind target antigens, but also facilitate the removal of target antigens. This could be a major advantage for developing more potent therapeutic antibodies.

The methodologies and technology discussed in this report was built by strategic genetic engineering of specific genes and modifier sequences to overcome current limitations such as complex synthesis and bioconjugation and enable production of therapeutic antibodies with naturally phosphorylated N-glycans from mammalian cell lines. Therefore, very similar upstream and downstream manufacturing processes can be utilized for producing PNCA antibodies as for current therapeutic antibodies.

We believe that the N-glycosylated peptide chimeric design with the S1S3 PTase co- expression could provide a viable path to produce recombinant therapeutic antibodies with natural phosphorylated N-glycans for targeted protein degradation in the lysosome as more effective treatments for infection, cancer, autoimmune diseases, and neurodegenerative diseases. We hypothesize that this technology can also be applied towards creating lysosome targeted antigen/ligand to remove problematic autoantibodies and developing new cell and gene therapies for long-term treatment.

## Acknowledgements

We thank members of M6P Therapeutics and Dr. Stuart Kornfeld lab for helpful discussions and support.

## Materials and Methods

### Construct and vector design

Protein sequences for antibody variable regions were obtained from anti-FLAG M2 (PDB ID: 2G60), anti-TNF adalimumab variable (US Patent US6258562B1), anti-EGFR cetuximab variable (PDB ID:1YY8) anti-PD-L1 atezolizumab variable (PDB ID:5X8L). Expression plasmids were generated by DNA fragments from Twist Bioscience and placed in pcDNA3.1, pTwist vector or variable regions ordered in pTwist CMV-hIgG1 or pTwist CMV-hIgk for expression of heavy or light chain respectively. PCR primers were from Azenta Biosciences. PCR amplifications using Q5 polymerase (NEB, M0491S) and restriction digest for ligation or HiFi cloning of fragments (NEB, M0492S). Large scale DNA purification was performed using a Zymo Maxi-prep kit (Zymo, D4203). The N-glycosylated peptide (NATGNSTSNSTGNSTSNSTGNATMMST) was designed based on NX(S/T) N-glycosylation sequence. Human CI-MPR cDNA (amino acids 1-2304) with C-tail HPC4 tag (EDQVDPRLIDGK) codon optimized for CHO expression was ordered from Genscript in pcDNA3.1 plasmid. All plasmid sequences were confirmed by Sanger or Full plasmid sequencing.

### Antibody production, purification and conjugation of Alexa Fluor-594

ExpiCHO cells (ThermoFisher, A29127) were cultured and transfected with plasmids for heavy chain and light chain expression with or without co-expression of S1S3 PTase plasmid following the manufacturer’s protocol (ThermoFisher, A29131). Cells were cultured for 7-8 days, and conditioned media was harvested and filtered with 0.22 um filter. Antibodies were purified from the conditioned media using Nab Protein G spin columns (ThermoFisher, 89957) and eluted in Pierce IgG Elution Buffer (ThermoFisher, 21004). Purified antibodies were dialyzed in PBS buffer with 1:300 – 500 ratios (sample volume to PBS) overnight, using slide-a-lyzer cassettes with 20 K molecular weight cut-off (ThermoFisher, 66005). 100 ug of purified antibodies were conjugated with Alexa Fluor- 594 using the Alexa Fluor 594 antibody labeling kit (ThermoFisher, A20185) following manufacturer’s protocol.

### Endoglycosidases analysis

5 uL of GFP conditioned medium or 1 ug of each purified antibodies was treated with PNGase-F (NEB, P0704L) or Endo-H (NEB, P0702L) following the manufacturing protocol in a total 20 uL reaction volume. 1 uL (50 ng) of sample was loaded for SDS-PAGE for western blot analysis.

### Cell culture

human fibroblast cells GM16648 (Coriell Institute for Medical Research) and Tay-Sachs patient fibroblast cells GM02968 (Coriell Institute for Medical Research) were cultured in MEM (Corning, 10-009-CV) with 15% FBS (Neuromics, FBS001). HepG2 (ATCC, HB-8065), Daoy (ATCC, HTB-186) and A431 (Sigma, 85090402) cells were cultured in MEM medium (Corning, 10-009-CV) with 10% FBS. MDA-MB-231 (Sigma, 92020424) and HEK293T (ATCC, CRL-3216) cells were cultured in DMEM (Corning, 10-013-CV) with 10% FBS.

### Generation of CI-MPR knockout cell line

HEK293T were transfected with pX330 containing sgRNA to target human IGF2R/CI-MPR using the oligos (caccg<u>AAGTGCAACCAGATCTCTCC</u> and aaacGGAGAGATCTGGTTGCACTTC) (Addgene plasmid ID: 42230) following the protocol [29]. Cells were selected with puromycin for 48 hours, and single cells plated in 96 well plates by dilution cells to 0.5 cell per 100 uL. Loss of CI-MPR expression was confirmed by Western blot.

### HexM uptake and activity

Cells were seeded in 12-well plate overnight. HexM- producing conditioned media (20 uL/well) alone or combined with antibody (20 nM, final concentration) with or without M6P (2 mM, final concentration) were added accordingly into each well for 4 hours. Cells were then harvested and washed with PBS twice. Cell pellets were lysed in M-PER buffer (ThermoFisher, 78501) with protease inhibitor (ThermoFisher,78440). HexM activity was detected by 4-methylumbelliferyl (4-MU) substrate method to α-Hexosaminidase activity (HexA). Briefly, 5 uL of cell lysate was incubated with 1 mM of 4-methylumbelliferyl 6-sulfo-N-acetyl-b-D-glucosaminide (Sigma, 454428) in 10 mM citrate phosphate buffer with 0.5% Triton-X-100, pH 4.2 for 1 hour. 4-MU fluorescence was measured by SpectraMax ID3 (Molecular Devices, Sunnyvale, CA) [30] with 360 nm excitation, 460 nm emission.

### Antibody internalization experiment

Human fibroblast cells (GM16648) were seeded in a 6-well plate with 3x10^5^ cells per well 24 hours before staining. 10-20 nM Alexa Fluor 594 labeled antibodies were mixed with the culture medium and applied to the cells for 6 hours. To co-stain with nuclei and lysosomes, medium containing 10 ug/ml of Hoechst 33342 stain (ThermoFisher, H3570) and 1 uM LysoTracker Green DND-26 (ThermoFisher, L7526) was loaded to cells for 10 minutes. Then cells were washed once with PBS buffer (Corning, 21-040-CV). Images were taken under the Invitrogen EVOS M5000 Imaging System with 20x objective. Data was further processed by Fiji Image J [31].

### mCherry-FLAG internalization experiment

1 x10^6^ HepG2 cells were plated in 12-well plate for 24 hours and then incubated with FLAG-mCherry (1 ug) with or without Ab- FLAG or PNCA-FLAG at concentrations of 20 nM for 24 h. Cells were washed three times with PBS buffer (Corning, 21-040-CV) before imaging and harvesting for western blotting analysis. Data was processed and quantified by Fiji Image J [31].

### TNFα internalization experiment

1 x10**^6^** HepG2 cells were plated in 12-well plate 24 hours ahead and incubated with TNFα alone or TNFα plus its antibody Ab-TNFα or PNCA-TNFα at the indicated concentrations for 24 h. Cells were washed three times and harvested and lysed in M-PER (ThermoFisher, 78501) with Protease and Phosphatases inhibitor cocktail (ThermoFisher, 78440) for western blot analysis.

### EGFR internalization experiment

5 x10^5^ Daoy cells were plated in 12-well plate and incubated with Ab-EGFR or PNCA-EGFR at various concentrations indicated for different amount of culture time as indicated. Cells were washed three times with PBS buffer and harvested and lysed in RIPA buffer (ThermoFisher 89900) with Protease and Phosphatases inhibitor cocktail (ThermoFisher, 78440).

### PD-L1 internalization experiment

5 x10^5^ Daoy or A431 cells were plated in 12-well plate and incubated with Ab-PD-L1 or PNCA-PD-L1 at various concentrations indicated for 24 h. Cells were washed three times with PBS buffer and harvested and lysed in RIPA buffer (ThermoFisher 89900) with Protease and Phosphatases inhibitor cocktail (ThermoFisher, 78440).

### SDS-PAGE and Western blot

Protein concentration was determined by BCA assay (ThermoFisher, 23227). 20 - 25 ug total protein were mixed with Pierce Lane Marker Reducing Sample Buffer (ThermoFisher, 39000) and boiled at 100 C for 5 minutes. Samples were loaded on Invitrogen NuPAGE 4-12% Bis-Tris Gels (ThermoFisher, NP0321BOX or NP0336BOX) and separated by 150V. Gels for Coomassie staining was performed by InstantBlue Coomasssie Stain (Abcam, ab119211). Gels for western were transferred to nitrocellulose membrane (Amersham, 10600011) overnight. After washing once with 1x PBST buffer (VWR, MSPP-IBB-171), membrane was blocked with 5% non-fat milk in PBST or 5% BSA (Sigma-Aldrich, A9085) at RT for 1 hour. Primary antibodies Rabbit anti-mCherry (ThermoFisher, PA5-34974) with 1:1000 dilution, Rabbit anti-TNFα (Cell Signaling, 6945s) with 1:1000 dilution, Rabbit anti-EGFR (Cell Signaling, 4267s) with 1:2000 dilution, Rabbit anti-PD-L1 (Abcam, ab228415) with 1:1000 dilution and Rabbit anti-GAPDH (ThermoFisher, PA1-987) with 1:2000 dilution in blocking buffer were incubated for another 1 hour at room temperature. HRP-linked anti-Rabbit IgG (Cell Signaling, 7074) was diluted with 1:5000 dilution for imaging under Azure 400. Data was quantified by Fiji Image J.

### Recombinant human CI-MPR production and purification

Human CI-MPR plasmid was transfected to ExpiCHO cells (ThermoFisher, A29127) following the manufacturer’s protocol (ThermoFisher, A29131). Cells were cultured for 7 days, and conditioned media was harvested and filtered with 0.22 um filter. The CI-MPR was purified by anti-HPC4 affinity agarose (Roche, 11815024001) following the manufacturer’s protocol.

### CI-MPR Affinity Chromatography

Pharmacia Fine Chemicals FPLC (Liquid Chromatography Controller LCC-500 and P-500 pumps) associated with 1 mL recombinant human CI-MPR agarose resin was used for the analysis. 3 ug of AF594 labeled antibody in 100 µL of buffer A (50 mM Imidazole, 150 mM NaCl, 2 mM EDTA, 5 mM beta-glycerophosphate, 0.05% Tx-100, 0.02% NaN3, pH 6.8) was injected and 0.5 mL/min flow rate was applied with total 5 mL buffer A to flow through any unbound material. 5 mL of linear gradient of 0 mM to 2 mM of mannose 6-phosphate (Sigma, M3655) by mixing buffer A and buffer B (50 mM NaOAc, 150 mM NaCl, 2 mM EDTA, 5 mM beta-glycerophosphate, 0.05% Tx-100, 0.02% NaN3, 10 mM M6P, pH 4.8) was applied to elute any loosely bound antibody. Final elution was performed with 5 mL buffer B to fully elute any bound material in the CI-MPR column. The column was then re-equilibrated with 4 volumes of buffer A. Every 0.2 mL fraction was collected in a clear 96-well plate during the process. Fraction collection plate was measured for AF594 fluorescence using the SpectraMax ID3 (Molecular Devices, Sunnyvale, CA) with 580 nm excitation, 620 nm emission. Raw fluorescence units were graphed against collection volume to determine the % antibody that binds to CI-MPR.

### CI-MPR plate binding assay

Purified bovine CI-MPR was immobilized on a clear flat bottom high binding 96-well plate (Costar, 3601) at a concentration of 1 ug/well. Following 2% bovine serum albumin (Sigma, A9085) blocking, samples were serially diluted into HEPES buffer (50 mM HEPES, 150 mM NaCl, 0.05% Tween-20, pH6.8) and incubated for one hour at room temperature on the CI-MPR coated plate. After incubation, the plate was washed three times with HEPES buffer and read for fluorescence using the SpectraMax ID3 (Molecular Devices, Sunnyvale, CA) with 580 nm excitation, 620 nm. Raw fluorescence units were graphed against the total concentration of antibody applied. The total fluorescence per dilution was also read to ensure antibodies at the same concentration yielded similar total fluorescence.

