## Supplementary material for "Targeted protein degradation in lysosome utilizing naturally produced bifunctional antibodies with high levels of mannose 6-phosphate glycans": suplemental figures

**Supplementary Figures**

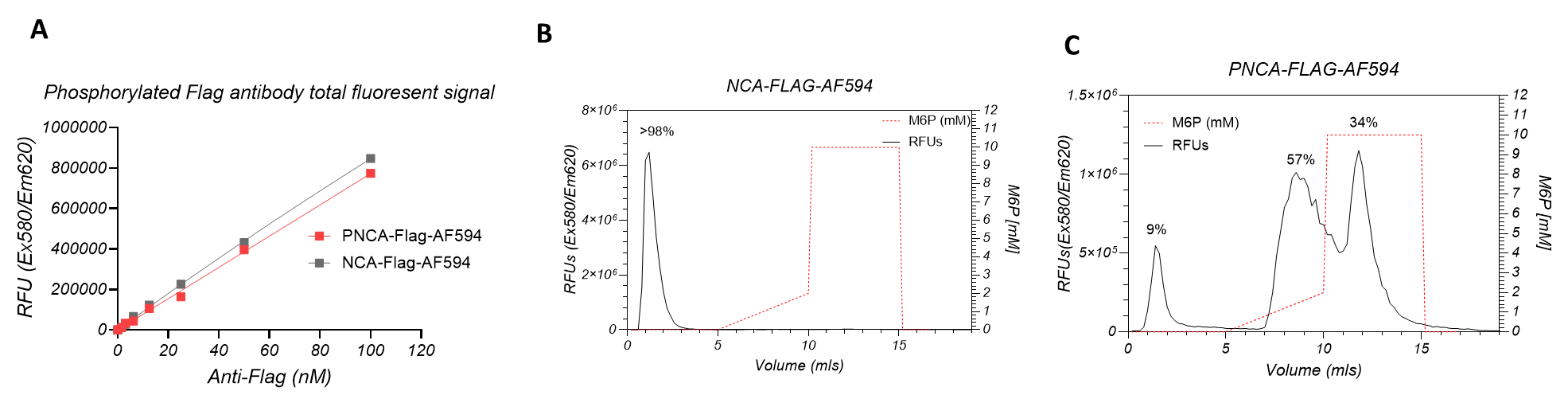

**Extended Data Figure 1.** A. Alexa fluor 594 (AF594) conjugated antibodies were analyzed for total fluorescence. PNCA-FLAG-AF594 (red) and NCA-FLAG-AF594 (grey) were diluted to several concentrations and fluorescence read in a plate reader at (Ex580/Em620). B. CI-MPR affinity chromatograph of NCA-FLAG-AF594. C. CI-MPR affinity chromatograph of PNCA-FLAG-AF594.

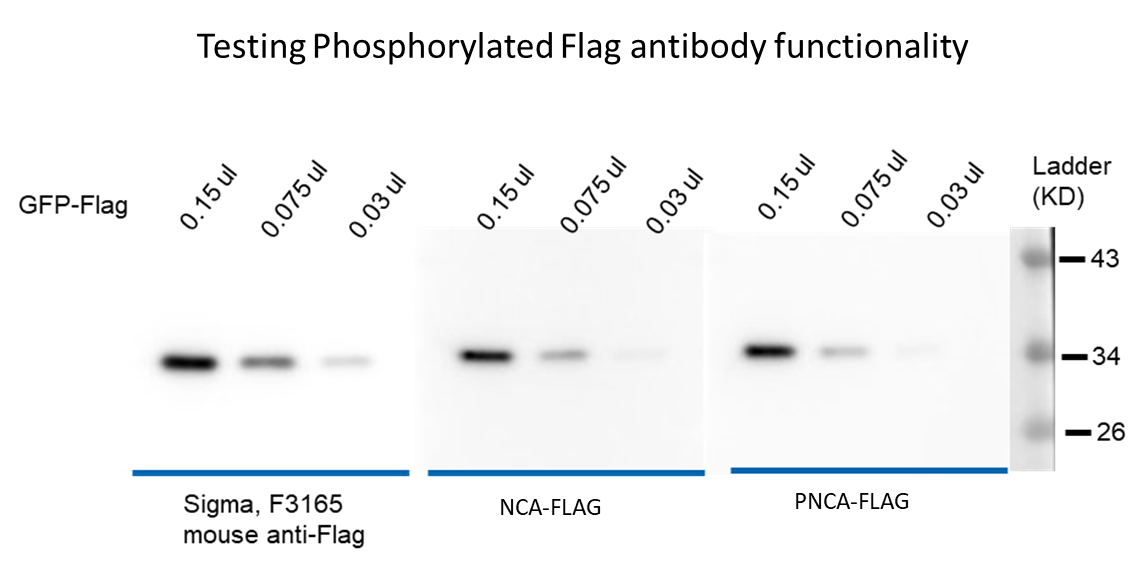

**Extended Data Figure 2.** Western blot analysis of antibody epitope recognition. Western blots were run with three dilutions of conditioned media containing soluble GFP-FLAG protein. Each blot was blotted with anti-FLAG antibodies (commercial Sigma F3165 mouse anti-FLAG, NCA-FLAG (M2) or PNCA-FLAG (M2) antibodies. Signal was detected with HRP-conjugated anti-mouse IgG.

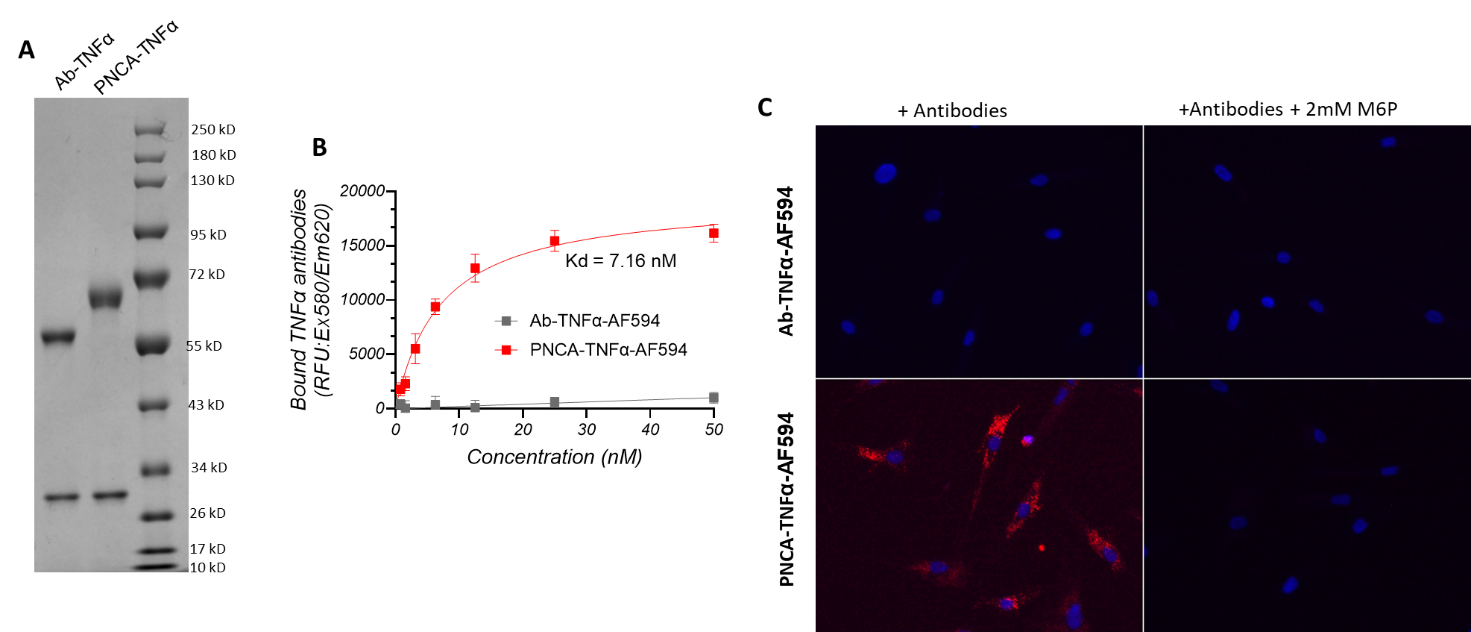

**Extended Data Figure 3.** A. Purified Ab-TNFα or PNCA-TNFα were analyzed by SDS-PAGE and Coomassie staining to visualize heavy and light chains. B. CI-MPR plate binding profile to examine the binding Kd (Kd=7.16 nM) for purified PNCA-TNFα-AF594 (red) and Ab-TNFα-AF594 (gray). C. AF594 conjugated antibodies (as in (B)) were incubated with human fibroblasts with or without 2 mM M6P (right panels). Uptake of antibody is indicated by red signal. Hoechst staining (blue) for nuclei.

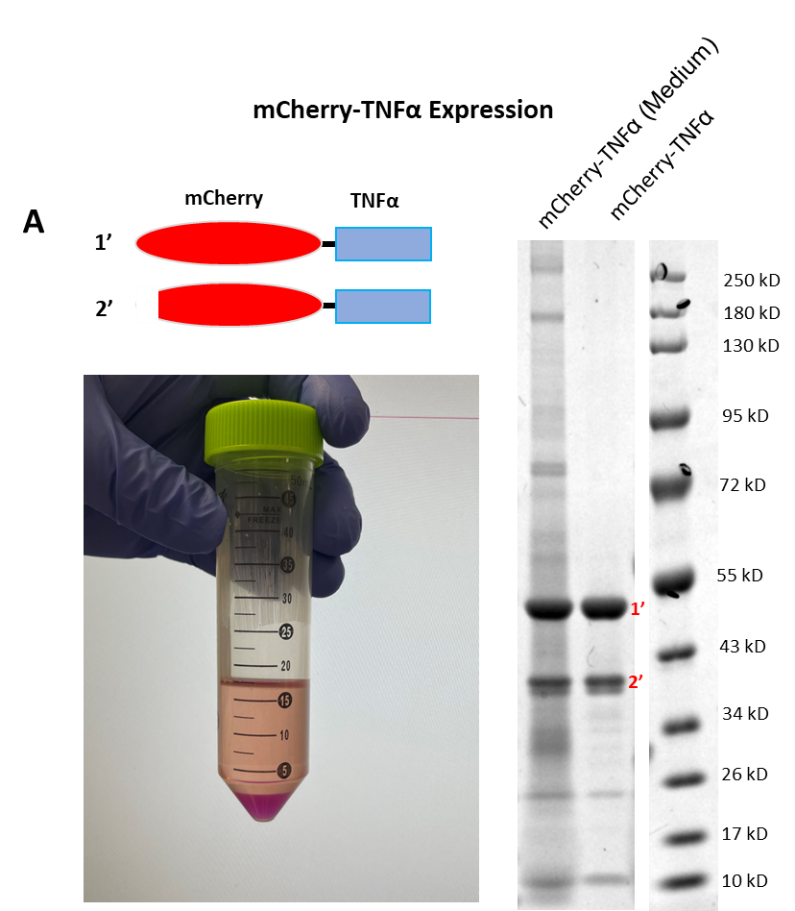

**Extended Data Figure 4.** (Top left) Cartoon of mCherry-TNFα fusion proteins. 1’ full amino acid sequence expressed as higher molecular weight band on Coomassie gel (right). 2’ truncated that runs as a lower molecular weight. Photo of mCherry protein expression in media and cell pellet (bottom left).

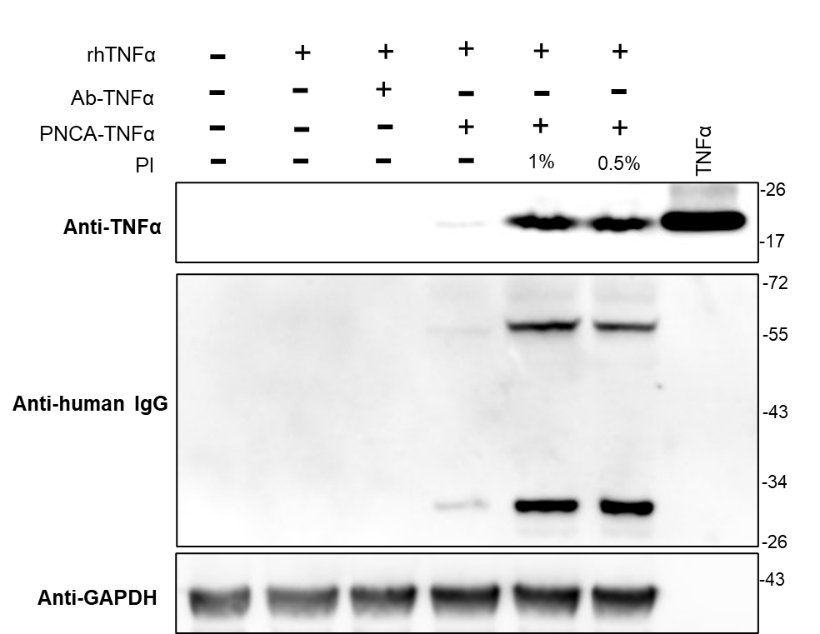

**Extended Data Figure 5.** HepG2 cells were incubated with recombinant TNFα (rhTNFα) and Ab-TNFα or PNCA-TNFα with or without protease inhibitors (PI). Cell lysates or 20 ng TNFα input were examined by Western blot. Blots were probed for TNFα, human IgG or GAPDH (loading control).

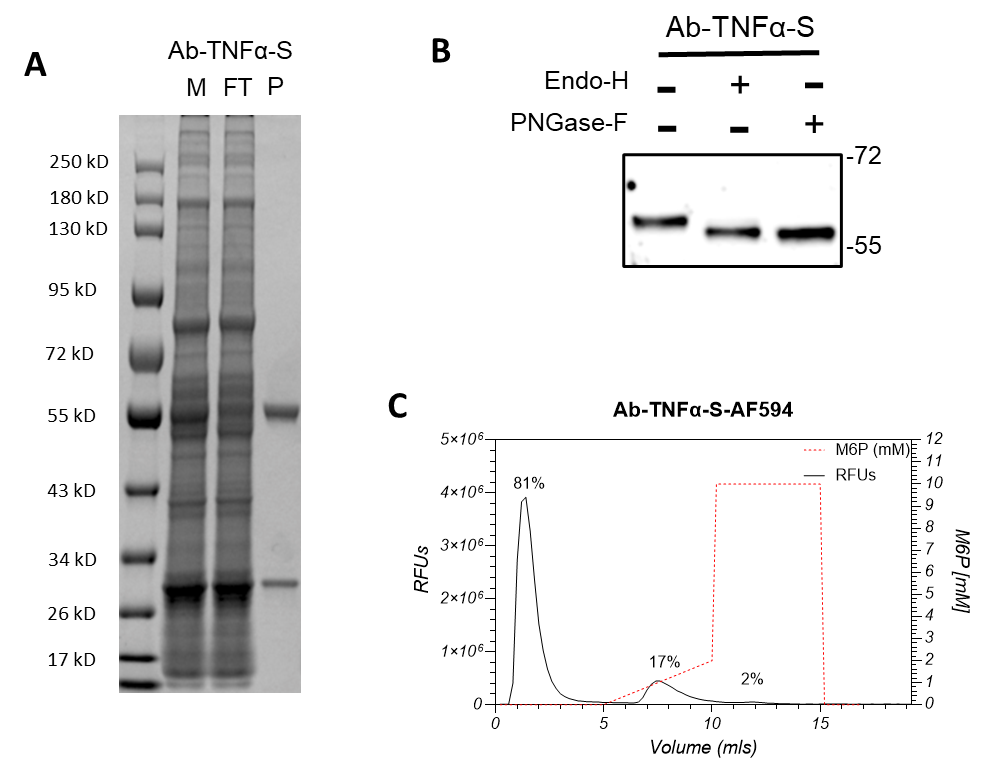

**Extended Data Figure 6.** A. Conditioned media and purified antibody produced by co-transfection of TNFα antibody sequence with S1S3 PTase (Ab-TNFα-S) were run on SDS-PAGE and examined by Coomassie stain. M: conditioned media; FT: flowthrough; P: purified protein. B. Purified Ab-TNFα-S as in (A) was treated with PNGase-F or Endo-H enzymes and analyzed by Western blot. (C) CI-MPR column binding profile for Ab-TNFα-S.

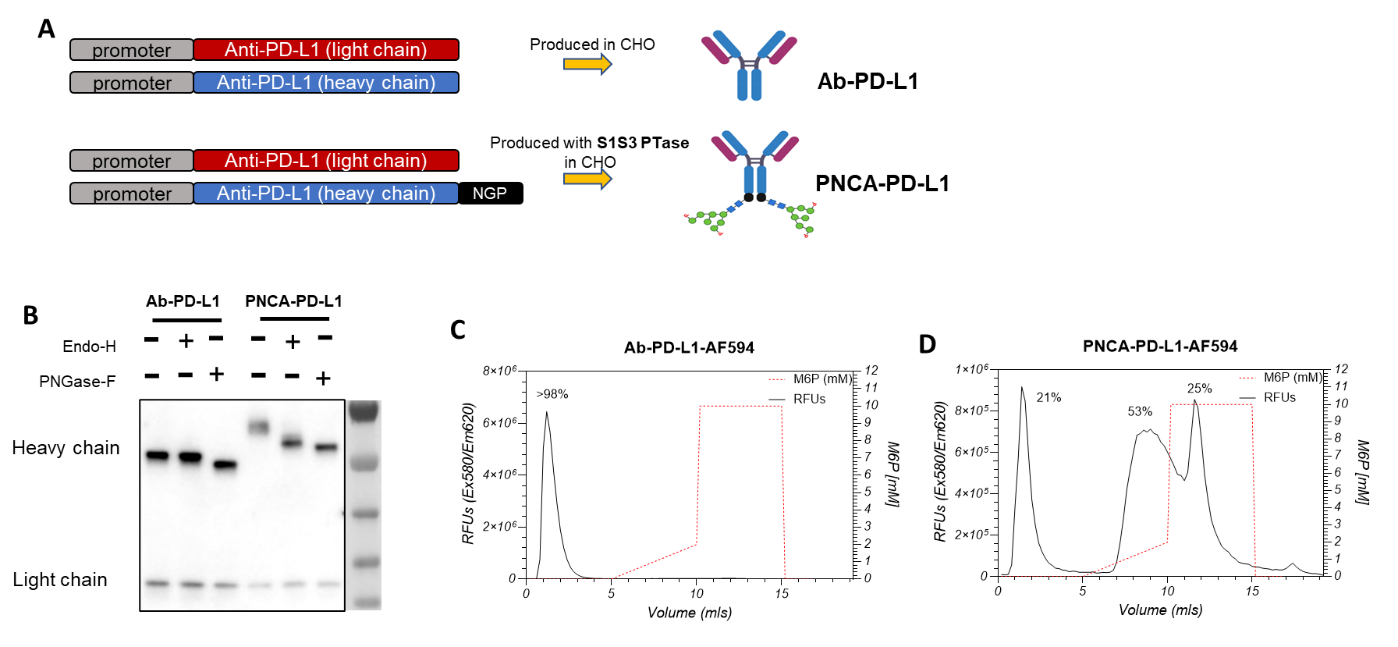

**Extended Data Figure 7.** A. Schematic for constructs expressed to produce Ab-PD-L1 by heavy and light chain constructs or PNCA-PD-L1 by co-expression of S1S3 PTase with heavy-NGP and light chain constructs. B. Purified Ab-PD-L1 or PNCA-PD-L1 were treated with PNGase-F or Endo-H enzymes and heavy and light chain examined by Western blot probed for human IgG. C. CI-MPR affinity chromatograph for purified AF594 conjugated Ab-PD-L1-AF594. D. CI-MPR affinity chromatograph for purified PNCA-PD-L1-AF594.

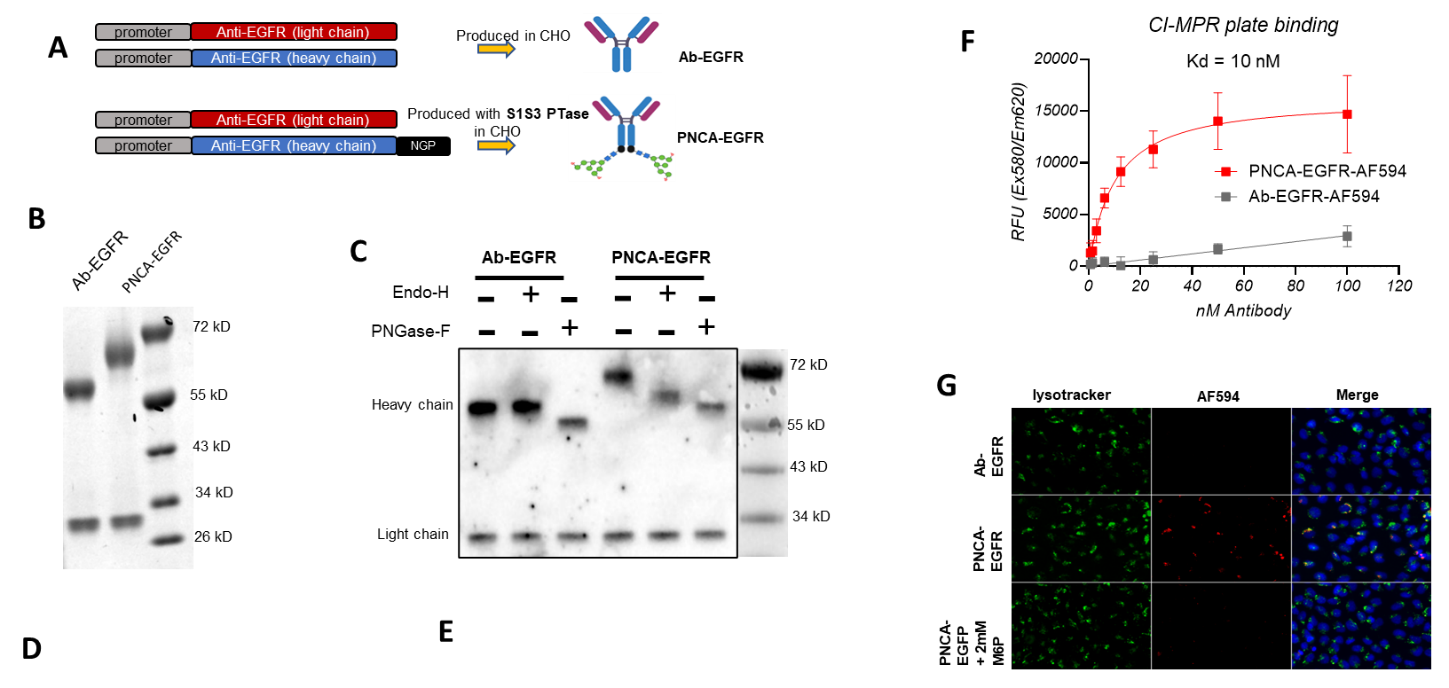

**Extended Data Figure 8.** A. Schematic for constructs expressed to produce Ab-EGFR by heavy and light chain constructs or PNCA-EGFR by co-expression of S1S3 PTase with heavy-NGP and light chain constructs. B. Purified Ab-EGFR and PNCA-EGFR were examined by SDS-PAGE and Coomassie staining. C. Purified Ab-EGFR and PNCA-EGFR as in (B) were treated with PNGase-F and Endo-H antibodies. Samples were examined by SDS-PAGE and Western blot probed for human IgG heavy and light chain. D. CI-MPR affinity chromatograph for Ab-EGFR-AF594. E. CI-MPR affinity chromatograph for PNCA-EGFR-AF594. F. Purified Ab-EGFR and PNCA-EGFR as in (B) were conjugated with alexa fluor 594 (AF594) and then examined for plate binding to CI-MPR to determine the binding affinity. Binding was detected by fluorescence for PNCA-EGFR-AF594 (red) or Ab-EGFR-AF594 (gray). G. HepG2 cells were incubated with AF594 conjugated purified Ab-EGFR-AF594 or PNCA-EGFR-AF594 and treated with or without 2 mM M6P. Antibody uptake is indicated by red signal. Lysosomes were stained by incubation with lysotracker (green), and nuclei were stained using Hoechst (blue).
